# Engineered type six secretion systems deliver active exogenous effectors and Cre recombinase

**DOI:** 10.1101/2021.05.11.443660

**Authors:** Steven J. Hersch, Linh Lam, Tao G. Dong

## Abstract

Genetic editing has revolutionized biotechnology but delivery of endonuclease genes as DNA can lead to aberrant integration or overexpression, leading to off-target effects. Here we develop a mechanism to deliver Cre recombinase as a protein by engineering the bacterial type six secretion system (T6SS). Using multiple T6SS fusion proteins, *Aeromonas dhakensis* or attenuated *Vibrio cholerae* donor strains, and a gain-of-function cassette for detecting Cre recombination, we demonstrate successful delivery of active Cre directly into recipient cells. Most efficient transfer was achieved using a truncated version of PAAR2 from *V. cholerae*, resulting in a relatively small (118 amino acid) ‘delivery tag’. We further demonstrate the versatility of this system by delivering an exogenous effector, TseC, enabling *V. cholerae* to kill *Pseudomonas aeruginosa*. This implicates that *P. aeruginosa* is naturally resistant to all native effectors of *V. cholerae* and that the TseC chaperone protein is not required for its activity. Moreover, it demonstrates that the engineered system can improve T6SS efficacy against specific pathogens, proposing future application in microbiome manipulation or as a next-generation antimicrobial. Inexpensive and easy to produce, this protein delivery system has many potential applications ranging from studying T6SS effectors to genetic editing.

**Importance:** Delivery of protein-based drugs, antigens, and gene-editing agents has broad applications. The type VI protein secretion system (T6SS) can target both bacteria and eukaryotic cells and deliver proteins of diverse size and function. Here we harness the T6SS to successfully deliver Cre recombinase to genetically edit bacteria without requiring the introduction of exogenous DNA into the recipient cells. This demonstrates a promising advantage over current genetic editing tools that require transformation or conjugation of DNA. The engineered secretion tag can also deliver a heterologous antimicrobial toxin that kills an otherwise unsusceptible pathogen, *Pseudomonas aeruginosa*. These results demonstrate the potential of T6SS-mediated delivery in areas including genome editing, killing drug-resistant pathogens, and studying toxin functions.

## Introduction

Genetic editing tools have provided incredible resources for DNA manipulation. In addition to the versatile and guided endonuclease, CRISPR-Cas9, other enzymes such as Cre recombinase transformed the field of biotechnology(1). Originally from the *Escherichia coli* P1 bacteriophage, Cre recombines genetic material to remove DNA flanked by *loxP* sites (‘floxed’), and this has been harnessed in diverse organisms ranging from bacteria to mice(1, 2). However, genetic editing enzymes have limitations, including that they must be introduced into a target cell. This is typically done by transferring DNA encoding the active enzyme, which can potentially cause unwanted mutations by integrating into the target chromosome or overexpressing the enzyme(3–5). Delivering genetic editors as proteins can potentially alleviate these issues by eliminating the possibility of DNA integration while simultaneously allowing for more control over enzyme dosage.

Bacteria encode numerous protein secretion systems such as the type 3 (T3SS) and type 4 secretion systems (T4SS), which are often virulence factors that pass unfolded effector proteins through a central pore into eukaryotic host cells(6). Several successful endeavors have demonstrated that these secretion systems can be harnessed for delivery of recombinant proteins(7, 8). Another, more recently discovered system, is the type six secretion system (T6SS)(9, 10). The T6SS resembles a molecular spear gun and uses physical puncturing to penetrate nearby bacteria or eukaryotic cells in order to deliver protein effectors with various destructive activities(11, 12). Importantly, the toxicity of the system towards prey cells is primarily driven by the effectors since the damage caused by the puncture itself appears negligible, as demonstrated by effectorless strains constructed in multiple species(13–17).

Rather than passing unfolded proteins through a pore like the T3SS and T4SS, the T6SS mounts effectors onto the spear structure, which permits folding prior to delivery. Effectors can be loaded in the long Hcp protein tube that is thrust forward, or mounted on a pointed spear-head comprised of a trimer of VgrG proteins and a sharp PAAR protein tip. Notably, effectors can be non-covalently attached for delivery (‘cargo’ effectors) or included as extended domains of the structural proteins, which are termed ‘evolved’ or ‘specialized’ effectors. Past attempts at delivering recombinant proteins using T6SSs have revealed key challenges(18) as well as some successes, which include delivery of β-lactamase into eukaryotic cells(19–21) and fusion of two exogenous effectors for secretion(21, 22). *Vibrio cholerae* encodes one of the best characterized T6SS and the T6SS has been particularly well characterized in strain V52, which is a non-pandemic strain with a constitutively active T6SS(9). *V. cholerae* V52 naturally employs five known effectors to inhibit macrophage or amoeba that phagocytose it, or to kill neighbouring bacteria including T6SS^+^ competitors such as *Aeromonas dhakensis*(23–26). In contrast, some species, such as *Pseudomonas aeruginosa*, survive attacks by the *V. cholerae* T6SS, but the mechanism of this resistance remains uncertain(27). The five T6SS effectors of *V. cholerae* include two evolved effectors, VgrG1 (containing an actin cross-linking domain) and VgrG3 (containing a lysozyme domain)(28, 29). The other three are cargo effectors, VasX, TseL and TseH, which are loaded onto VgrG2, VgrG1 or PAAR2, respectively(17, 30, 31). Furthermore, the V52 strain is genetically malleable and was previously attenuated by deletion of three non-T6SS toxins, *rtxA*, *hlyA*, and *hapA*(9), rendering it less toxic towards eukaryotic cells. This makes it a potentially viable donor for downstream applications of protein delivery into eukaryotic cells or in the context of a host organism.

In this work we demonstrate that the T6SS can be harnessed by engineering fusion tags that facilitate delivery of active Cre recombinase directly into neighbouring recipient cells. We picked Cre recombinase as the delivered exogenous protein for the following reasons:

1. Cre has previously been used to demonstrate engineered T3SS-mediated delivery(32).
2. The size of Cre (343 amino acids) is comparable to the effector domains of VgrG1 and VgG3.
3. Cre instigates efficient and specific DNA recombination allowing for detection in recipient cells.
4. Cre/*loxP* recombination is commonly used in numerous species and therefore Cre delivery has immediate potential applications.
5. Cre delivery represents a proof-of-principle for transfer of active genetic editing enzymes into the cytoplasm of target cells, which establishes the system for future development using targeted endonucleases such as CRISPR-Cas9.

We develop a relatively small (118 amino acids), yet highly efficient, delivery tag that achieves substantial levels of *loxP*-site recombination in recipients; this highlights its potential for future genetic editing applications. We further demonstrate the utility of the system by efficiently delivering an exogenous T6SS effector that grants *V. cholerae* the ability to kill *P. aeruginosa*, demonstrating the application of the system towards studying effector activity or targeting particular bacterial prey.

## Results

### Cre fused to VgrG is secreted in a T6SS-dependent manner

We hypothesized that evolved VgrG proteins would be most amenable to secreting exogenous proteins due to their natural fusion to effector domains. Specifically, by replacing the effector domain with a protein of interest, toxic effector activity is removed while leaving the natural linker to the VgrG domain intact, which may be important for proper folding and loading onto the T6SS tip. Since *V. cholerae* V52 has two evolved VgrGs and one of the best studied T6SSs, we employed it to deliver Cre recombinase cargo in initial T6SS engineering experiments.

To generate VgrG-Cre fusions, we first inserted *vgrG* genes – truncated prior to their effector domains – into a plasmid vector with a C-terminal 3x V5 tag (Fig. 1A); these effectorless VgrG plasmids served as ‘no-Cre’ controls. Notably, the remaining linker region of VgrG1 includes a proline-rich region (PRR; codons 686-725) where 35% of the residues are proline, including six in a row. VgrG3 also has a linker between the effector domain and the VgrG domain; the ‘conserved domains analysis feature’ of BLAST(33) revealed that this region could be split into a linker associated with the lysozyme domain (646-717) and another linker region (540-645). Since it was unclear if these linkers were important for T6SS-mediated secretion, we introduced Cre recombinase at two different locations: either maintaining the entire linker (VgrG1_725_-Cre_3V5_ and VgrG3_717_-Cre_3V5_) or removing the PRR or lysozyme-associated linker domains (VgrG1_685_-Cre_3V5_ and VgrG3_645_-Cre_3V5_, respectively)(Fig. 1A).

**Figure 1:**
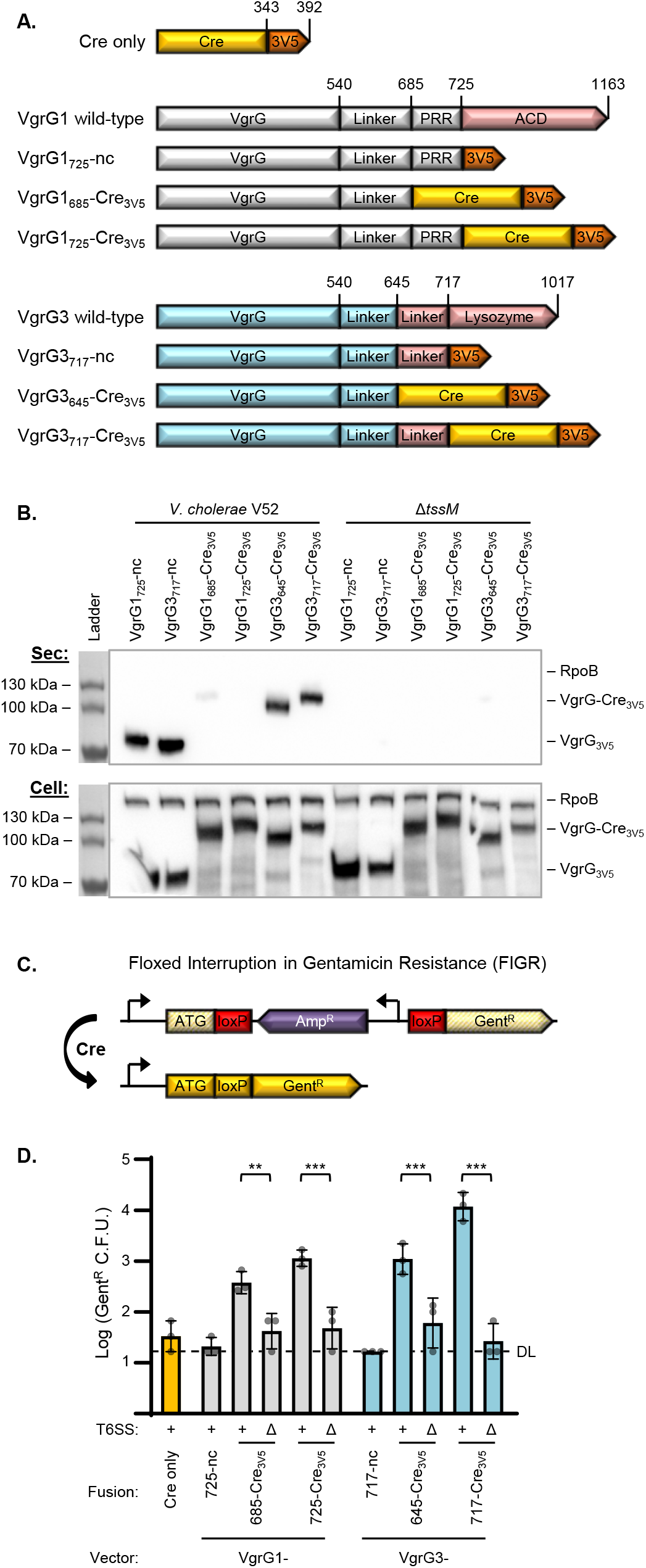
Active VgrG-Cre fusions are secreted and delivered to recipient cells in a T6SS-dependent manner. **A.** Depiction (not to scale) showing fusions of Cre recombinase (yellow) and 3x V5 tag (orange) to VgrG1 and VgrG3 of *V. cholerae* V52. VgrG1 (grey) wild-type includes a proline-rich region (PRR) and an actin cross-linking domain (red; ACD). VgrG3 (blue) wild-type includes a lysozyme domain (red) with a linker region leading up to the peptidoglycan binding domain beginning at codon 718. nc, ‘no Cre’. **B.** Western blot showing secretion of VgrG1 and VgrG3 fusions from wild-type *V. cholerae* V52 but not an equivalent *tssM* mutant. ‘Sec’ (top) shows secreted fractions, ‘Cell’ (bottom) shows cell pellet lysates. In addition to α-V5, α-RpoB antibody was included as a cytoplasm control. Representative of three independent replicates. **C.** Depiction of the ‘Floxed Interruption in Gentamicin Resistance’ (FIGR) cassette. An ampicillin resistance cassette (purple) – encoded in the reverse direction and flanked by loxP sites (red) – interrupts the open reading frame of a gentamicin resistance gene (yellow striped). Exposure to Cre removes the block and allows expression of the gentamicin resistance gene (yellow). Promoters are shown as 90° arrows. Figure is not to scale. ATG, Gent^R^ start codon. **D.** Recovery of *V. cholerae* Gent^R^ colony forming units (indicative of Cre-mediated FIGR cassette recombination) after delivery from wild-type (+) or Δ*tssM* (Δ) *V. cholerae* with indicated Cre fusions to VgrG vectors. DL, detection limit. One-way ANOVA with Sidak’s multiple comparison test comparing wild-type and Δ*tssM* donors with equivalent delivery fusions. Recovery from Δ*tssM* samples were not significantly above ‘Cre only’ or ‘no Cre (nc)’ controls. **, p<0.01; ***, p<0.001.

We electroporated the VgrG-Cre plasmids into the *V. cholerae* V52 strain and tested secretion by western blotting for the V5 antigen. Both ‘no-Cre’ controls were secreted from wild-type bacteria but not from an equivalent T6SS-inactive (Δ*tssM*) mutant (Fig. 1B). This data suggests T6SS-dependent release of these truncated VgrGs since the proteins were fully expressed in the Δ*tssM* mutant (whole-cell fractions) and no cell lysis was observed (measured as RNA polymerase, RpoB, in the secreted fractions). Moreover, both VgrG3-Cre fusions exhibited prominent T6SS-dependent secretion (Fig. 1B).

In contrast, the VgrG1-Cre fusions were only slightly detectable in the secreted fractions (Fig. 1B). This suggests that VgrG1-Cre is either unstable in the extracellular milieu or is secreted much less than VgrG3-Cre. To test if VgrG1-Cre was secreted at all, we tested its ability to deliver the cargo effector, TseL. Normally TseL binds to VgrG1 for delivery and a V52 *vgrG1* mutant does not kill a sensitive strain lacking the TseL immunity gene, *tsiV1*(30, 31). The VgrG1_725_-Cre_3V5_ fusion was able to complement TseL delivery in a Δ*vgrG1* strain (Fig. S1), suggesting that VgrG1_725_-Cre_3V5_ can bind to TseL and be secreted into neighbouring cells. We therefore continued to examine the VgrG1-Cre fusions in further assays.

### Active VgrG-Cre fusions are delivered into recipient cells

We next sought to determine if the engineered fusions could deliver active Cre recombinase into neighbouring cells. We constructed a plasmid-based cassette for detecting Cre-mediated recombination in recipient cells: The plasmid contains a floxed (flanked by *loxP* sites) ampicillin resistance (Amp^R^) cassette that interrupts translation of a gentamicin resistance (Gent^R^) gene (Fig. 1C). We termed this plasmid pFIGR for <u>f</u>loxed <u>i</u>nterruption in <u>g</u>entamicin <u>r</u>esistance. Before Cre-recombination, pFIGR grants ampicillin resistance but the cells remain gentamicin sensitive. Upon Cre-mediated recombination, the block is lifted and the Gent^R^ gene is expressed, granting gentamicin resistance to the bacteria. This provides a simple method for detecting Cre recombination as growth of recipient cells on media containing gentamicin. Notably, recipients would only become sensitive to ampicillin if all copies of pFIGR (approximately 20 / cell) lost the Amp^R^ cassette.

To avoid killing the recipients with T6SS effectors, we used the same *V. cholerae* V52 background – encoding all immunity genes – as the recipient. Furthermore, to prevent re-secretion of the fusions by the T6SS of recipient cells, the Δ*tssM* mutant was used as the recipient strain. Finally, for enumeration of total recipients, we also introduced a kanamycin resistant (Kan^R^) plasmid that is compatible with pFIGR.

After combining donor cells and recipients containing pFIGR, we found that all four of the VgrG-Cre fusions led to significant levels of Gent^R^ colony forming units (C.F.U.) (Fig. 1D). The VgrG3_717_-Cre_3V5_ construct was the most effective, leading to about 500-fold more Gent^R^ C.F.U. than controls. Importantly, we confirmed that the gain of gentamicin resistance required Cre, since controls lacking Cre demonstrated background levels of Gent^R^ C.F.U.. Moreover, three pieces of evidence support that the observed gentamicin resistance was dependent on direct T6SS-mediated delivery: 1) a Cre-only control that lacks a VgrG ‘delivery tag’ did not increase the number of Gent^R^ colonies. 2) Wild-type donors demonstrated significantly more Cre delivery (Gent^R^ C.F.U.) than T6SS-null (Δ*tssM*) strains (Fig. 1D, S2A). 3) Attempts to deliver Cre under conditions that are prohibitive of the T6SS, including in liquid media or with a barrier separating the donor and recipient cells, failed to increase the yield of Gent^R^ colonies (Fig. S2B, C). Notably, Cre delivery does not appear to recombine every copy of pFIGR in a cell, since Amp^R^ (non-recombined FIGR) and Kan^R^ (total recipient) C.F.U. counts were indistinguishable – suggesting that some copies of pFIGR retain the Amp^R^ gene (Fig. S2A).

Since the T6SS has been associated with horizontal gene transfer, we also considered that transfer of either pFIGR or the Cre donor plasmid (which is chloramphenicol resistant; Cm^R^) could result in a strain containing both plasmids – leading to Gent^R^ colonies. However, this did not appear to occur since the Cre-only control plasmid did not increase the numbers of Gent^R^ C.F.U. (Fig. 1D) and not a single Cm^R^Gent^R^ double-resistant colony was detected across all replicates (data not shown). These data suggest that the observed Gent^R^ colonies resulted from genuine T6SS-mediated delivery of active Cre protein.

Data from this initial Cre delivery experiment revealed some limitations to be addressed: First, the VgrG-Cre fusions appear to restrict the donor cells, leading to reduced recovery compared to no-Cre or Cre-only controls (Fig. S2D). This was independent of T6SS activity, suggesting that the VgrG-Cre fusions result in undetermined toxicity or may exert a resource strain on donor cells expressing the large recombinant proteins. The second limitation was that we observed reduced recipient survival with wildtype donors compared to Δ*tssM* (Fig. S2A), suggesting that T6SS effectors were killing recipient cells despite the recipient strains encoding all immunity genes. These limitations are addressed in the following sections.

### Using a PAAR2-Cre delivery vector improves donor fitness

Since expressing the VgrG-Cre fusions conveyed a fitness cost on the donor cells (Fig. S2D; as described above), we sought an alternative delivery fusion that would not inhibit the donor cells. PAAR2 in *V. cholerae* does not have an effector domain but, in addition to its N-terminal PAAR domain, it does include a C-terminal tail for recruiting the effector, TseH(17). Full-length PAAR2 is only 176 amino acids long, which is a significantly smaller ‘T6S-tag’ than the VgrG fusions; we hypothesized that the smaller size would reduce the metabolic burden of expressing it. We fused Cre to either full-length PAAR2 (PAAR2_FL_) or a truncated version (PAAR2_106_) that has only 12 amino acids of the C-terminal tail (Fig. 2A). Notably, since the VgrG1-Cre plasmid was used as a template, the cloning method left a 12 amino acid remnant of VgrG1 (codons 714-725, downstream of polyproline region); these residues act as a further linker between PAAR2 and Cre.

**Figure 2:**
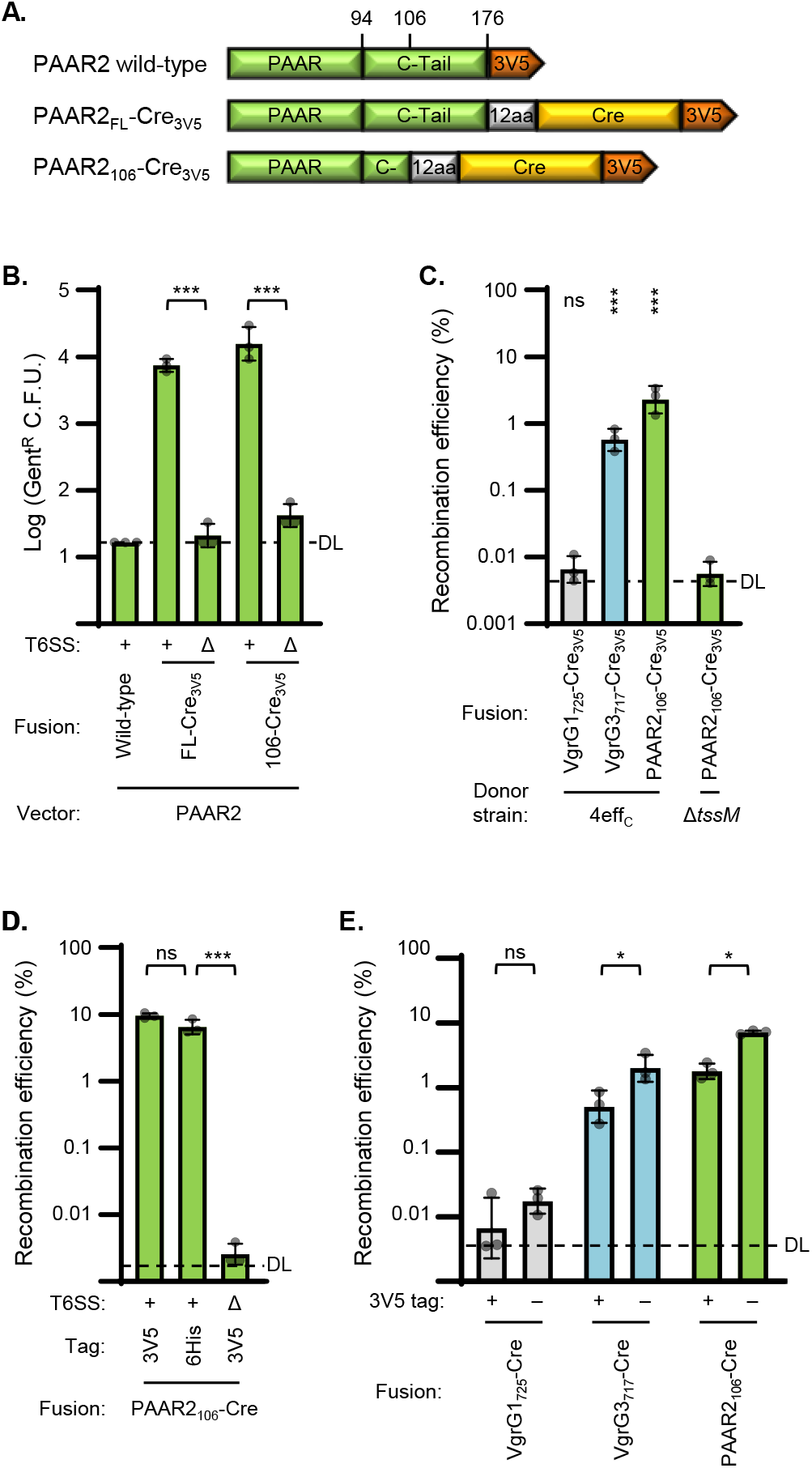
Improving Cre delivery using PAAR2 fusions, an effectorless donor strain, and removing 3V5 tags. **A.** Depiction (not to scale) showing fusions of Cre recombinase (yellow) and 3x V5 tag (orange) to PAAR2 (green) of *V. cholerae* V52. Fusions include the full-length C-terminal tail (C-Tail) of PAAR2 (FL) or just 12 amino acids of it (C-); 12 amino acids (12aa) are also leftover from the VgrG1_725_-Cre_3V5_ construct (codons 714-725) used as template for PAAR2 insertion. **B.** Recovery of Cre-recombined recipient bacteria (Gent^R^ C.F.U.) after delivery from wild-type (+) or Δ*tssM* (Δ) *V. cholerae* with indicated Cre fusions to PAAR2 vectors. DL, detection limit. One-way ANOVA with Sidak’s multiple comparison test: Recovery from Δ*tssM* samples were not significantly above no-Cre (PAAR2 wild-type) control. ***, p<0.001. **C.** Cre fusion delivery from *V. cholerae* with catalytically inactivated antibacterial effectors (4eff_C_) or Δ*tssM* (Δ). Data shows recombination efficiency (Gent^R^ / Kan^R^ C.F.U. recovered; recombined / total recipients) as a percentage. DL, approximate detection limit. One-way ANOVA with Dunnett’s multiple comparison test comparing each sample to the Δ*tssM* donor strain. ***, p<0.001; ns, not significant. **D.** Recombination efficiency after delivery from 4eff_C_ (+) or Δ*tssM* (Δ) *V. cholerae* encoding PAAR_106_-Cre with either a 3V5 or 6His tag. DL, approximate detection limit. One-way ANOVA with Tukey’s multiple comparison test. ***, p<0.001; ns, not significant. **E.** Recombination efficiency after delivery from 4eff_C_ strain encoding indicated Cre fusions with the 3V5 tag present (+) or removed (−). DL, approximate detection limit. One-way ANOVA with Sidak’s multiple comparison test comparing samples with and without 3V5 tags. *, p<0.05; ns, not significant.

Delivery of either PAAR2_FL_-Cre_3V5_ or PAAR2_106_-Cre_3V5_ led to significant and substantial Cre-mediated recombination in recipient cells (Fig. 2B). The number of Gent^R^ recipients following PAAR2_106_-Cre_3V5_ delivery was slightly higher than PAAR2_FL_-Cre_3V5_, and comparable to the most effective fusion tested previously, VgrG3_717_-Cre_3V5_ (Fig. 2B compared to Fig. 1D). The no-Cre control (wild-type PAAR2) and all Δ*tssM* donors yielded undetectable or background amounts of recombination, demonstrating that recombination in recipients required T6SS-mediated secretion of Cre. As expected, delivery of the PAAR2-Cre fusions was also contact-dependent, since delivery in liquid media or with a barrier separating the donor and recipient cells yielded negligible Gent^R^ C.F.U. (Fig. S3A, B). Importantly, expression of the PAAR2-Cre fusions did not reduce donor cell fitness (Fig. S3C), suggesting that they did not cause a fitness cost like the VgrG-Cre fusions. However, delivery of these fusions from wild-type donor cells still demonstrated some toxicity to the recipients (Fig. S3D).

### Inactivating donor effectors restores recipient survival

Despite using *V. cholerae* recipients – with all immunity genes intact – we consistently observed some degree of toxicity towards the recipients that was independent of which Cre fusion was being delivered (Fig. S2A, S3D). To address this limitation, we switched donor strains to a previously generated V52 strain that has all four of its antibacterial effectors catalytically inactivated (4eff_C_)(16, 17). The 4eff_C_ strain showed no toxicity to recipients (Fig. S4A). Notably, delivery of VgrG-Cre fusions was reduced when delivered from the 4eff_C_ strain compared to the wild-type donor (Fig. S4B compared to Fig. 1D). However, delivery of the PAAR2_106_-Cre_3V5_ fusion was similar from the 4eff_C_ donor strain as from the wild-type (Fig. S4B compared to Fig. 2B). These data support using the 4eff_C_ strain to solve the recipient toxicity issue.

### Measuring recombination efficiency and further improving efficacy

Using the 4eff_C_ donor strain, toxicity was prevented and total recipient recovery was consistent regardless of which fusion construct was delivered (Fig. S4A). This allowed for measuring recombination efficiency by dividing the number of recombined recipients (Gent^R^ C.F.U.) by the number of total recipients (Kan^R^ C.F.U.). Using this metric, we observed up to about 2.5% recombination efficiency using the PAAR2_106_-Cre_3V5_ construct (Fig. 2C).

We considered that the C-terminal 3x V5 tag might inhibit delivery or Cre activity; we experimented using a 6x His tag (6His) or removing the detection tag entirely. The 6His tag did not reduce recombination efficiency (Fig. 2D), suggesting that 6His-tagged proteins can be successfully secreted using the PAAR2_106_ vector. Removal of the detection tag significantly improved recombination efficiency for both VgrG3_717_- and PAAR2_106_-Cre fusions (Fig. 2E), raising the efficiency to over 7%. Oddly, in the 6His experiment (Fig. 2D), the PAAR2_106_-Cre_3V5_ construct exhibited up to 10% recombination efficiency, highlighting that there is some variability between experiments.

### T6SS effectors can serve as delivery vectors

Up to this point we have demonstrated delivery of Cre as a fusion to T6SS structural genes, VgrG1, VgrG3 and PAAR2 of *V. cholerae*. To further determine the versatility of T6SS-mediated protein delivery, we fused Cre to the C-termini of the three *V. cholerae* cargo effectors, VasX, TseL and TseH (Fig. 3A). The VasX-Cre construct did not facilitate recombination in recipient cells (Fig 3B) or kill VasX-sensitive prey (Fig. S5), suggesting that this fusion was inactive or failed to be secreted. However, both TseL and TseH fusions demonstrated significant Cre delivery (Fig. 3B). The efficiency of TseH-Cre delivery rivalled VgrG_717_-Cre_3V5_ but was lower than PAAR2_106_ fusions (Fig. 3B compared to Fig. 2E). These findings demonstrate that non-structural proteins can also serve as T6SS delivery tags to deliver customized cargo proteins.

**Figure 3:**
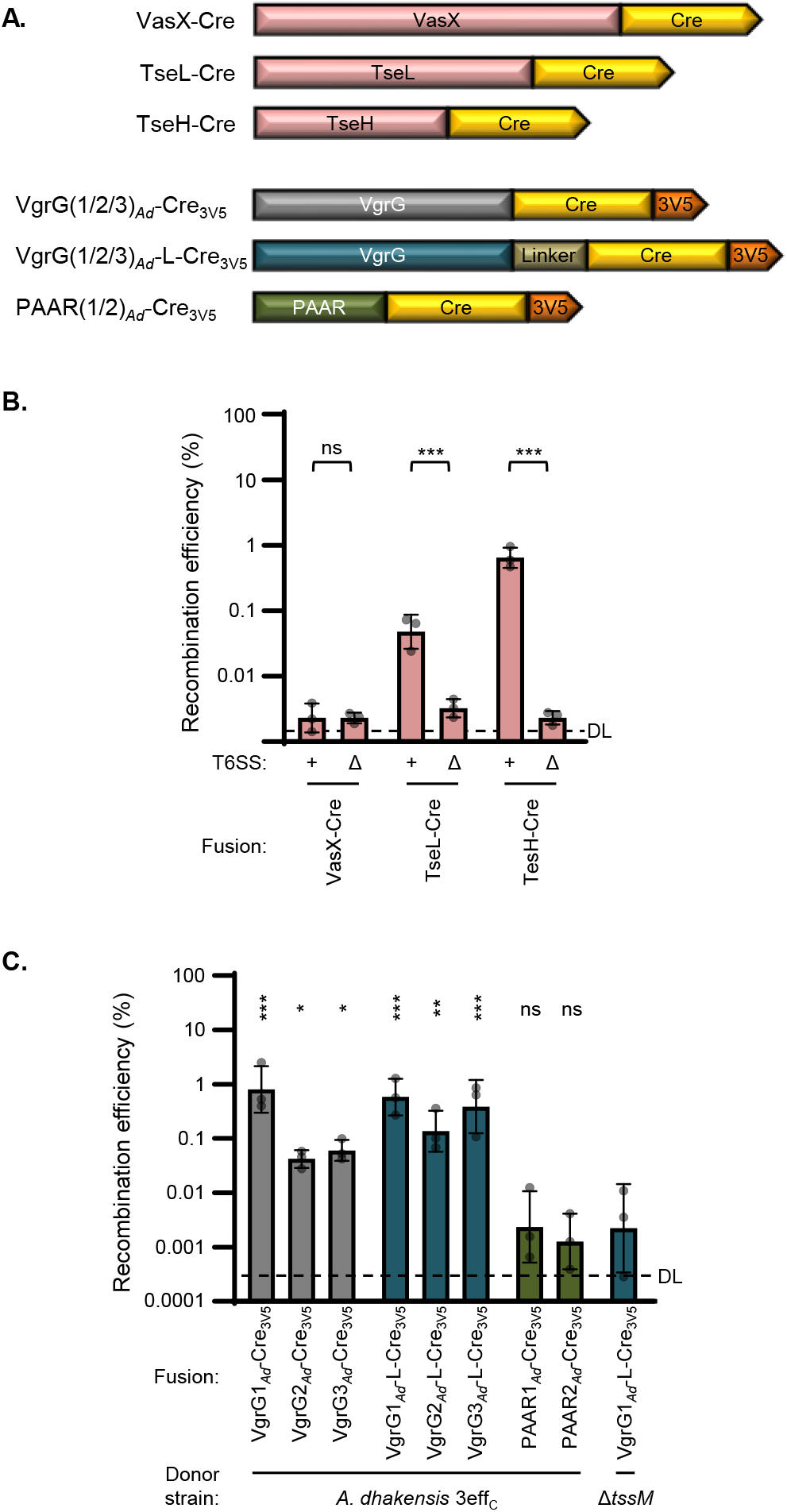
Active Cre can be delivered as effector fusions and by *A. dhakensis*. **A.** Depiction (not to scale) showing fusions of Cre recombinase (yellow) and 3x V5 tag (orange) to *V. cholerae* T6SS effectors (red) or to *A. dhakensis* PAAR proteins (dark green) and VgrG proteins with (dark blue) or without (dark grey) a flexible linker inserted. **B.** Recombination efficiency after delivery from *V. cholerae* 4eff_C_ (+) or Δ*tssM* (Δ) strains encoding Cre fusions to indicated effector proteins. DL, approximate detection limit. One-way ANOVA with Sidak’s multiple comparison test comparing 4eff_C_ and Δ*tssM* donors with equivalent delivery fusions. ***, p<0.001; ns, not significant. **C.** Recombination efficiency after delivery of Cre fusions from *A. dhakensis* with catalytically inactivated antibacterial effectors (3eff_C_) or Δ*tssM* (Δ). DL, approximate detection limit. One-way ANOVA with Dunnett’s multiple comparison test comparing each sample to the Δ*tssM* donor. *, p<0.05; **, p<0.01; ***, p<0.001; ns, not significant.

#### *A. dhakensis* is another proficient donor strain

We were curious if other species could also be employed for T6SS-mediated delivery of recombinant proteins. We tested another aquatic species that has a constitutively active T6SS, *A. dhakensis* strain SSU(30, 34). We fused Cre to each of the three VgrGs and two PAAR proteins present in *A. dhakensis* (Fig. 3A). We then introduced these constructs into an *A. dhakensis* strain with its three antibacterial effectors, TseI, TseP and TseC, all catalytically inactivated (3eff_C_)(35). This strain showed no toxicity towards *V. cholerae* recipients (Fig. S6B)(35).

All three VgrG fusions led to significant Cre delivery from the T6SS^+^ donor compared to an equivalent Δ*tssM* strain (Fig. 3C); delivery of VgrG1*_Ad_*-Cre_3V5_ yielded the highest recombination efficiency. In contrast, neither of the *A. dhakensis* PAAR fusions demonstrated signs of delivery. Since none of the *A. dhakensis* VgrG proteins have effector domains, we attempted to better mimic an evolved VgrG by introducing a flexible linker between Cre and the VgrG domain (Fig. 3A). The linker appeared to improve delivery for VgrG2*_Ad_* and VgrG3*_Ad_*, but not VgrG1*_Ad_* (Fig. 3C). Notably, none of the fusion constructs appeared to inhibit donor fitness and the 3eff_C_ strain showed no recipient toxicity compared to the Δ*tssM* strain (Fig. S6). Cumulatively, these findings demonstrate that the T6SS of multiple species can be engineered to deliver active Cre recombinase. Additionally, this transfer can cross species barriers (*A. dhakensis* delivery to *V. cholerae* recipients) if the donor’s antibacterial effectors are inactivated.

We also examined delivery to a recipient strain with a chromosomal copy of FIGR instead of the pFIGR plasmid (approximately 20 copies per cell). Delivery of the VgrG3_Ad_-L-Cre_3V5_ fusion from *A. dhakensis* resulted in slight but significant levels of recombination efficiency (Fig. S7). This suggests that chromosomal FIGR can detect Cre-mediated recombination to a measurable level, though it appears to be less efficient than the plasmid-borne cassette.

### The PAAR2_106_ T6S tag enables *V. cholerae* to deliver a recombinant effector to kill *P. aeruginosa*

To examine if engineered T6S tags can deliver proteins other than Cre, we focused on delivering exogenous T6SS effectors. Customizing effector delivery could be used to study effector activities or to maximize efficacy against particular prey. As a proof-of-principle, we employed *V. cholerae* and *P. aeruginosa*. It was previously shown that the T6SS of *V. cholerae* does not kill *P. aeruginosa*(27), however the mechanism remains unclear. One possibility is that *V. cholerae* effectors are not active against *P. aeruginosa*; alternatively, the *V. cholerae* T6SS might not be able to puncture the *P. aeruginosa* cell envelope to deliver the effectors(17, 27, 36).

We employed T6S fusions to elucidate this mystery: We noticed that *A. dhakensis* can use its T6SS to kill *P. aeruginosa* and this lethality is abolished when the three antibacterial effectors are catalytically inactivating (Fig. 4A). By inactivating two effectors at a time – leaving a single active effector in the strain – we determined that TseP and TseC were effective against *P. aeruginosa* and TseC was more consistent (Fig. 4A). In light of this, we replaced Cre (of the PAAR2_106_-Cre fusion) with TseC and its downstream immunity gene (*tsiC*) from *A. dhakensis*. Both wild-type and the 4eff_C_ strain of *V. cholerae* were able to deliver this PAAR2_106_-TseC*_Ad_* fusion, resulting in greatly reduced survival of competing *V. cholerae* (Fig. 4B). Moreover, expressing PAAR2_106_-TseC*_Ad_* enabled both wild-type and 4eff_C_ *V. cholerae* to kill *P. aeruginosa* (Fig. 4C). Importantly, this was T6SS- and TseC*_Ad_*-dependent, suggesting that reduced *P. aeruginosa* survival results from T6SS-mediated delivery of the *A. dhakensis* effector from *V. cholerae*. Accordingly, *V. cholerae* delivering PAAR2_106_-TseC*_Ad_* inhibited *P. aeruginosa* to a similar degree as the *A. dhakensis* strain with TseC as its only active effector (Fig. 4A compared to C).

**Figure 4:**
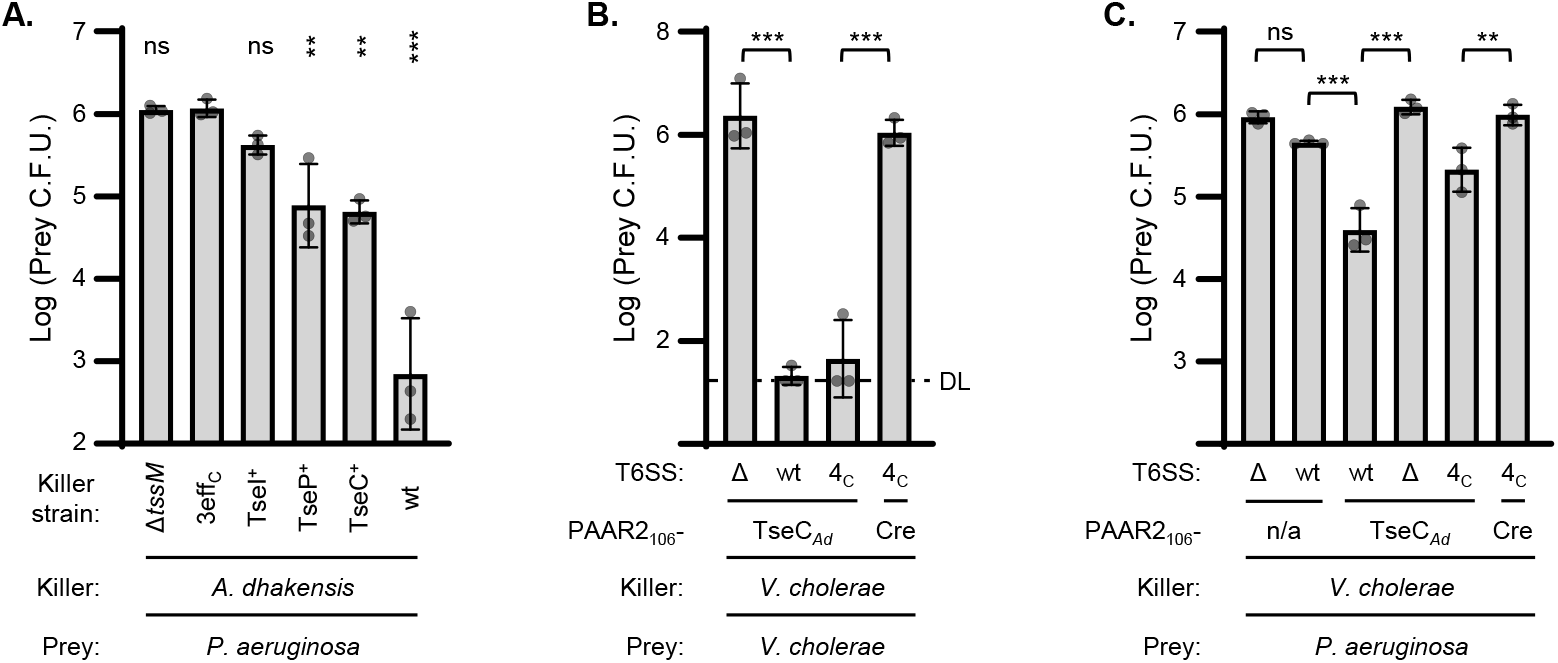
Fusion of PAAR2_106_ to the *A. dhakensis* effector, TseC, empowers *V. cholerae* to kill *P. aeruginosa*. **A.** Prey *P. aeruginosa* recovery after incubation with *A. dhakensis* killer cells. Prey *P. aeruginosa* has Δ*hsiB* (Δ*tssB*) mutations in all three T6SS. Killer strains have T6SS-null mutation (Δ*tssM)* or a wild-type T6SS with all effectors intact (wt), all antibacterial effectors catalytically inactivated (3eff_C_), or a single active effector (indicated) while the other effectors are inactivated. One-way ANOVA with Dunnett’s multiple comparison test comparing all samples to the 3eff_C_ killer strain. **, p<0.01; ***, p<0.001; ns, not significant. **B.** Prey *V. cholerae* (Δ*tssM*) recovery after incubation with *V. cholerae* killer cells encoding PAAR2_106_ fusions to *A. dhakensis* TseC (TseC*_Ad_*) or Cre as a control. Killer strains have Δ*tssM* (Δ) or a wild-type T6SS with all effectors intact (wt) or all native antibacterial effectors catalytically inactivated (4eff_C_). DL, detection limit. One-way ANOVA with Sidak’s multiple comparison test. ***, p<0.001; ns, not significant. **C.** Prey *P. aeruginosa* (Δ*hsiB* mutations in all three T6SS) recovery after incubation with *V. cholerae* killer cells with no fusion construct (n/a) or encoding PAAR2_106_ fusions to *A. dhakensis* TseC (TseC*_Ad_*) or Cre as a control. Killer strains have Δ*tssM* (Δ) or a wild-type T6SS with all effectors intact (wt) or all native antibacterial effectors catalytically inactivated (4eff_C_). One-way ANOVA with Sidak’s multiple comparison test. **, p<0.01; ***, p<0.001; ns, not significant.

## Discussion

In this work we developed multiple fusion tags allowing the T6SSs of *V. cholerae* and *A. dhakensis* to efficiently deliver the targeted endonuclease, Cre, or an exogenous effector, TseC*_Ad_*, directly into neighbouring cells (Fig. 5). We also showed activity of the system with a 6His-tagged protein, potentially leading to a downstream method for lysis-free purification of secreted fusion proteins. Additionally, we constructed a novel gain-of-function Cre detection cassette (FIGR) and identified the *A. dhakensis* T6SS effectors that are most active against *P. aeruginosa*.

**Figure 5:**
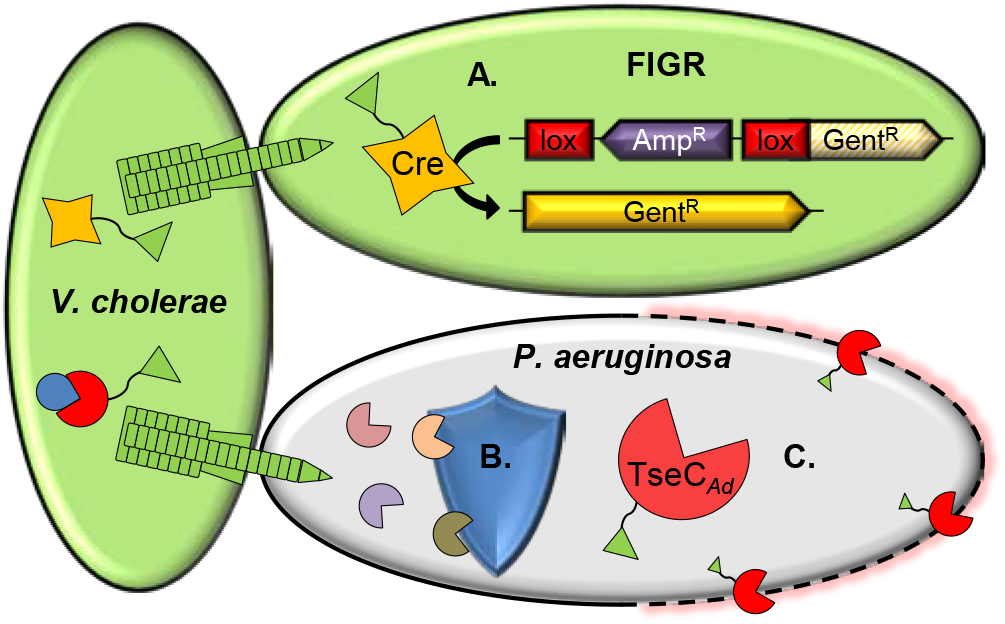
Model of engineered T6SS-mediated delivery of Cre recombinase and TseC_*Ad*_. **A.** Engineered T6SSs of *V. cholerae* or *A. dhakensis* donor cells deliver active Cre recombinase protein. Delivered Cre instigates recombination of the floxed interruption in gentamicin resistance (FIGR) cassette in recipient cells. **B.** *P. aeruginosa* exhibits natural resistance (by unknown mechanisms) to all four antibacterial effectors of wild-type *V. cholerae.* **C.** *V. cholerae* engineered to deliver the *A. dhakensis* effector, TseC*_Ad_*, gains the ability to kill *P. aeruginosa*.

The delivery fusion results described in this work reveal details of T6SS function. For instance, since Cre was able to access the DNA of recipient cells, it implicates that the T6SS tip can puncture into the cytoplasm. This supports previous works using nuclease effectors, the reuse of secreted T6SS components in recipient sister cells, and the finding that VgrG3 of *V. cholerae* has evolved to re-export from the cytoplasm of recipient cells to access its peptidoglycan target in the periplasm(22, 37). Another finding was that Cre could be delivered as a fusion to PAAR2 of *V. cholerae* but not when fused to either PAAR protein of *A. dhakensis*. Likely, this results from the extended C-terminal domain of PAAR2 that is not present in the other PAAR proteins. This region acts as a linker (even when truncated in the PAAR2_106_ construct) that potentially improves the folding or functionality of Cre or the PAAR domain. This suggests that effectors that are attached to PAAR domains must include a sufficient linker or else impede folding or functionality.

We employed the PAAR2_106_ T6S tag to engineer *V. cholerae* to deliver the *A. dhakensis* effector, TseC. This led to a number of significant findings: 1) TseC was fully active when delivered from *V. cholerae*, suggesting that TseC’s chaperone (TEC) protein is not required for proper folding or activity(30). However, *V. cholerae* also encodes two DUF4123 TEC proteins, so it remains possible that they can act in lieu of the *A. dhakensis* chaperone(30). 2) Since the *tseC* fusion gene demonstrated full activity after switching to a new host organism and delivery mechanism, this provides further empirical evidence that domain swapping can facilitate new effector integration. This supports previous works examining the evolution of the T6SS by horizontal gene transfer and the modular nature of effectors(22, 38–41).

The findings additionally highlight the evolutionary advantage of accumulating exogenous effectors since *V. cholerae* equipped with PAAR2_106_-TseC devastated neighbouring *V. cholerae* populations (Fig. 4B). Moreover, it gained the ability to kill *P. aeruginosa*, which suggests that the *V. cholerae* T6SS is capable of delivering effectors into *P. aeruginosa* but these prey cells have the ability to resist killing by all four of *V. cholerae*’s natural antibacterial T6SS effectors. This resistance may not be complete, as delivery of *V. cholerae* effectors can instigate a retaliatory ‘tit-for-tat’ response in *P. aeruginosa*(17, 27). Additionally, delivering PAAR2_106_-TseC*_Ad_* from the wild-type donor appeared to kill *P. aeruginosa* more than delivery from the 4eff_C_ strain, suggesting that there may be some level of synergy between TseC and the *V. cholerae* effectors. Nonetheless, the *V. cholerae* effectors alone did not significantly reduce *P. aeruginosa* survival. The mechanism of this resistance remains to be determined in future work but could involve cross-protection by *P. aeruginosa*’s array of immunity genes or alternative tolerance systems such as stress response-mediated damage repair. This finding further supports a growing body of work highlighting that certain effectors are more efficacious against particular prey cells or under particular conditions(16, 17, 42–44).

The antibacterial activity of the T6SS can be harnessed to target specific species for manipulating microbiomes or as a potential next-generation antimicrobial(21, 22, 45). To achieve this, it is crucial to be able to equip the T6SS with a customized selection of effectors. Effector discovery used to be a major challenge of the field due to sequence divergence, but multiple approaches have been developed to overcome this earlier challenge(23, 30, 46–48). Now the pool of divergent effectors comprises thousands of proteins in hundreds of species, including many that are genetically intractable. Thus, the ability to swap an effector into a different T6SS delivery species will allow for further study of effectors – and defences against them – in isolation from their native delivery strains. The fusions described here establish delivery mechanisms for auxiliary effectors to maximize efficacy against particular target species, thereby enhancing the potential of the T6SS for future applications.

Finally, the engineered T6SS provides advantages over existing platforms for delivery of genetic editing enzymes as proteins: 1) The T6SS can deliver directly into the target cell cytoplasm without requiring a receptor. This grants an advantage over secreted proteins, which require receptors for uptake, deliver only to the periplasm of bacteria, or both. 2) The T6SS can deliver ready-folded proteins, providing an advantage over T3SSs or T4SSs for cargo that is inactivated by denaturation. 3) Due to its receptor-independent mechanism, the T6SS has a broad spectrum of target cells including both eukaryotic and prokaryotic cells. The system described here provides the first example of delivering a genetic editing enzyme as a protein into bacteria.

Throughout the process of optimizing the T6SS delivery fusions, we employed Cre recombinase. Notably, Cre acts as a tetramer, requiring at least four molecules to be delivered to recipients before recombination can occur. Since three VgrG proteins are delivered per firing event, the VgrG-Cre fusions potentially deliver multiple Cre molecules at once. However, it is not known if the VgrG trimer dissociates in recipient cells or if Cre instigates recombination while still assembled in a VgrG trimer. In contrast, the PAAR_106_ fusion tag can only deliver one protein per T6SS firing event but may promote Cre functionality due to the smaller T6S tag domain. Because it functions as a tetramer, Cre recombination likely understates delivery efficiency, which may increase with the use of monomeric enzymes in the future.

In addition to the immediate potential applications of Cre delivery, it primarily acts as a proof-of-principle: The successful delivery of Cre – and subsequent recombination in recipient cells – marks an important landmark for the use of the T6SS in genetic engineering methods. Vector integration remains a significant constraint in genetic editing, but protein delivery using the T6SS can potentially eliminate this issue. Dosage can also be readily regulated to limit off-target effects by repressing T6S tag expression or using antibiotics to rapidly end endonuclease delivery by killing the donor bacteria. Future work, including with additional gene editing tools such as CRISPR-Cas9, will further develop this protein delivery system.

## Materials and Methods

### Bacterial strains and growth conditions

Strains used in this study are listed in Supplementary Table S1A. All *V. cholerae* were from the V52 strain background with deletions of *rtxA*, *hlyA*, and *hapA* (RHH)(9). Unless otherwise indicated, recipients were *V. cholerae* Δ*tssM* with pFIGR and pBAD33k as an independent, compatible plasmid for total C.F.U. enumeration. *P. aeruginosa* prey were strain PAO1 with deletion of all three *tssB* (*hsiB*) genes to prevent T6SS retaliation(49), and included the pPSV37 plasmid for selection. Bacteria were grown shaking at 37 °C in LB media (0.5% NaCl) or on LB agar plates. For plasmid maintenance and selection of recipient cells for C.F.U. counts, antibiotics were added to final concentrations of 2.5 μg/mL chloramphenicol, 100 μg/mL carbenicillin, 50 μg/mL kanamycin, 20 μg/mL gentamicin.

### Plasmid construction

Plasmids used in this study are listed in Supplementary Table S1B. Plasmids were generated using Gibson cloning(50) or overlapping PCR mutagenesis(51) and verified by Sanger sequencing. In brief, pFIGR was generated by first inserting *loxP* sites on both sides of the ampicillin resistance cassette of pBAD24. This floxed Amp^R^ gene was then inserted (in the reverse-direction) before the second codon of the gentamicin resistance gene in pPSV37. For chromosomal FIGR, a version of FIGR with Kan^R^ instead of Amp^R^ (FIGR_kan_) was transferred into the pGP-Tn7 plasmid allowing for transposon-mediated insertion (with the pSTNSK helper plasmid) at the chromosomal *attTn7* site(52).

Delivery fusions were generated in the pBAD33 plasmid backbone (arabinose inducible, chloramphenicol resistant) with a C-terminal 3x V5 tag. Truncated VgrG proteins lacking effector domains were amplified from gDNA by PCR and inserted into the vector, followed by insertion of Cre recombinase at sites indicated in Fig. 1A. PAAR2-Cre fusions were generated by replacing VgrG1 with PAAR2_FL_ or PAAR2_106_, leaving a 12 amino acid remnant of VgrG1 as a linker between PAAR2 and Cre. Subsequent effector, *A. dhakensis*, and TseC fusions were generated by Gibson cloning to replace VgrG or Cre in existing plasmids.

### Secretion assay and western blot

Secretion assays were conducted similar to described previously(16). In brief, strains were grown for 3 hours with arabinose added to 0.4% to induce fusion gene expression. Cultures were centrifuged and pellets stored for sonication as cell lysate samples. Meanwhile supernatants were passed through 0.22 μm filters, trichloroacetic acid (TCA) solution was added to final concentration of 20%, and placed at −20 °C overnight. Supernatants were then centrifuged at 15,000 g for 20 min at 4 °C, pellets washed with 100% acetone, and the resultant pellet resuspended in SDS-loading dye for analysis by western blotting:

As described previously(15, 53), proteins were resolved by polyacrylamide gel electrophoresis, transferred to a nitrocellulose membrane, blocked with 5% skim milk in TBST buffer (50 mM Tris, 150 mM NaCl, and 0.05% Tween 20, pH 7.6) for 1 h at room temperature, then incubated with primary antibodies overnight at 4 °C. Blots were washed 3 times in TBST, incubated with anti-mouse HRP-conjugated secondary antibody (Cell Signaling Technology) for 1 h, then detected using ECL solution (Bio-Rad). Monoclonal antibodies to the V5 epitope tag and RpoB, the beta subunit of RNA polymerase used as a cytoplasmic control, were purchased from GeneTex and NeoClone, respectively.

### T6SS competition and delivery assays

T6SS activity assays were conducted similar to described previously with minor modifications(17): ‘Donor’ or ‘killer’ strains were subcultured in LB (with antibiotic for plasmid maintenance as needed) for 3 hours with arabinose added to 0.4% (0.01% for *A. dhakensis*) for the last hour. ‘Recipient’ or ‘prey’ strains were from overnight cultures. For wild-type Cre donors and equivalent controls, donor and recipient were mixed at a 5:1 ratio. Competition assays and donor experiments with *V. cholerae* 4eff_C_ (or *A. dhakensis* 3eff_C_) donors employed 10:1 ratios. Mixtures were spotted on LB plates containing 0.1% arabinose (0.01% for *A. dhakensis* donor strains) and incubated for 3 hours at 37 °C. Agar plugs containing the mixed bacteria were removed using wide-bore pipette tips, resuspended in PBS, serially diluted and plated for colony forming units (C.F.U.) on LB plates containing antibiotics as indicated in figure axes and legends. Data are reported as either log C.F.U. recovered on particular antibiotic plates, or as ‘Recombination Efficiency’ calculated as Gent^R^ C.F.U. / Kan^R^ C.F.U.. If no colonies were observed across triplicate plating, samples were listed as the ‘detection limit’ represented as if 0.5 colonies were counted. For recombination efficiency, ‘detection limit’ line is approximate since the denominator (total C.F.U.) varies between samples.

## Acknowledgements

This work was supported by grants from Canadian Institutes of Health Research (CIHR) and Canadian Natural Sciences and Engineering Research Council (NSERC) to T.G.D.. T.G.D. is also supported by a Government of Canada Research Chair award, and a Canadian Foundation for Innovation grant (CFI-JELF). L.L. was supported by an Alberta Innovates Health Solutions (AIHS) Graduate Student Scholarship, and S.J.H. holds a CIHR Postdoctoral Fellowship.

## Author contributions

S.J.H. and T.G.D. conceived the project, S.J.H. and L.L. generated fusion constructs, S.J.H. designed, performed and analysed experiments, as well as prepared the original manuscript and figures.

T.G.D. supervised the study and guided manuscript revision.

## Supplemental Material legends

**Supplemental Figure S1:**
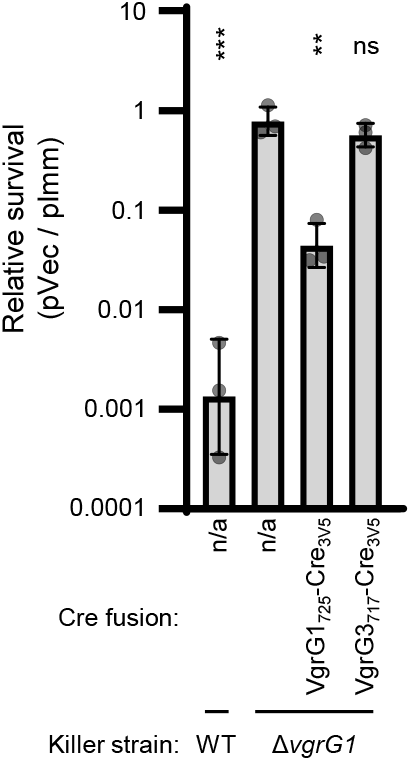
The VgrG1_725_-Cre_3V5_ fusion can deliver TseL to kill sensitive prey. T6SS competition assay showing killing by wild-type V52 (WT), or a V52 strain lacking the *vgrG1* gene (Δ*vgrG1*) and complemented with no plasmid (n/a), VgrG1_725_-Cre_3V5,_ or VgrG3_717_-Cre_3V5_. Prey are a V52 strain lacking the TseL immunity gene, *tsiV1*, and complemented with either empty vector (pVec) or *tsiV1* on a plasmid (pImm). Survival of the prey is shown as a ratio of recovered C.F.U. (pVec / pImm). One-way ANOVA with Dunnett’s multiple comparison test comparing each killer strain to Δ*vgrG1* with no plasmid. **, p<0.01; ***, p<0.001; ns, not significant.

**Supplemental Figure S2:**
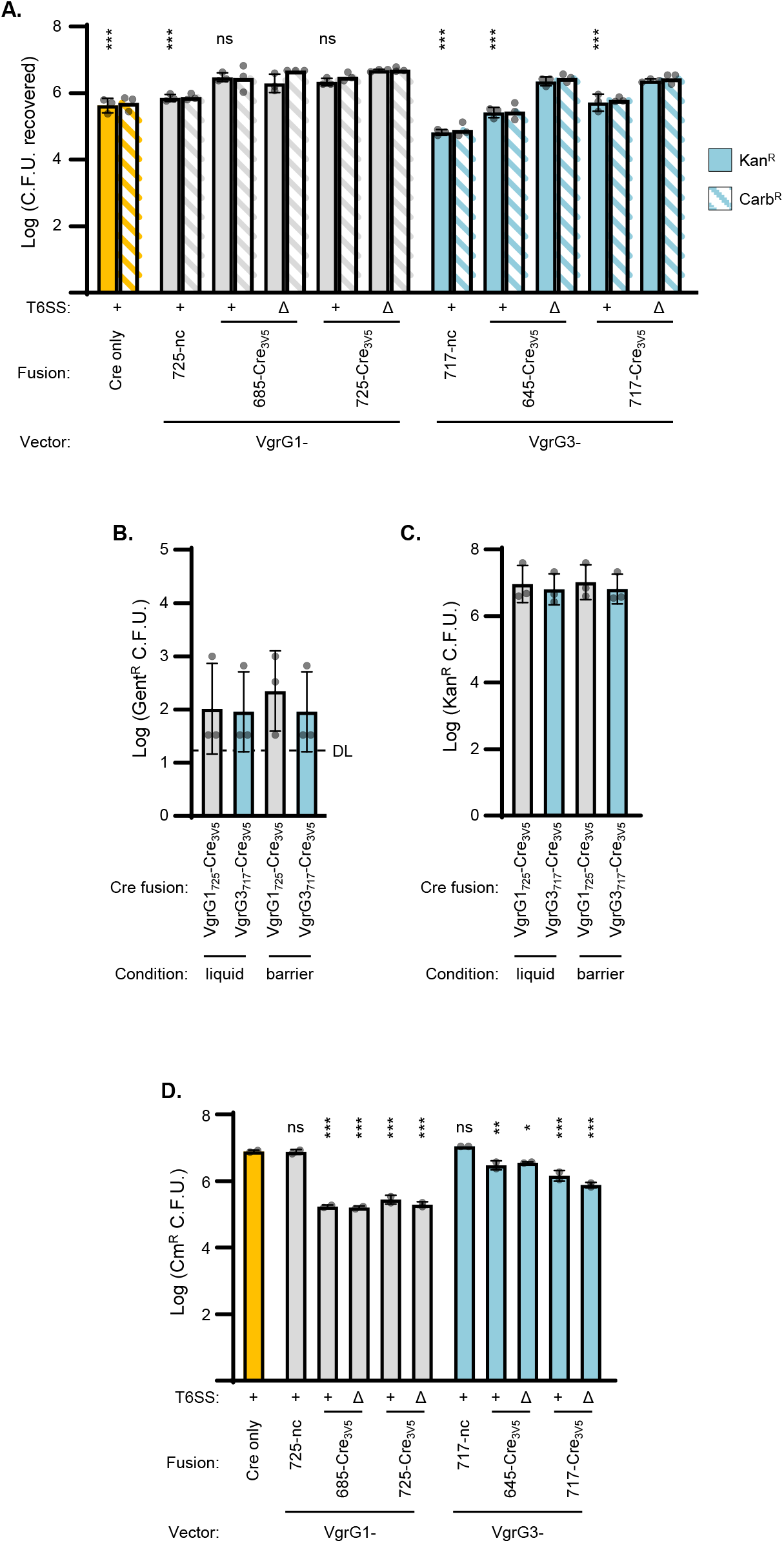
VgrG fusions total donor and recipient survival, liquid or barrier controls, related to Figure 1D. **A.** Recovery of *V. cholerae* Kan^R^ (total recipients) or Carb^R^ (recipients with at least one non-recombined copy of FIGR) colony forming units after delivery from wild-type (+) or Δ*tssM* (Δ) *V. cholerae* with indicated Cre fusions to VgrG vectors. One-way ANOVA with Sidak’s multiple comparison test comparing wild-type and Δ*tssM* donors with comparable delivery fusions. Kan^R^ statistics are shown, similar results were obtained for Carb^R^. No significant differences comparing Kan^R^ and Carb^R^ results for any delivery fusion strains. ***, p<0.001; ns, not significant. **B, C.** Cre delivery in liquid suspension (liquid) or when donor and recipient were separated by a nitrocellulose membrane (barrier). Recovery of *V. cholerae* colony forming units after incubating with wild-type *V. cholerae* with indicated Cre fusions. One-way ANOVA analyses indicate no significant differences. **B.** Gent^R^ C.F.U. indicative of Cre-mediated FIGR cassette recombination. DL, detection limit. **C.** Total recipient C.F.U. (Kan^R^). **D.** Recovery of *V. cholerae* Cm^R^ colony forming units (total donor cell recovery) with indicated Cre fusions to VgrG vectors after incubation with *V. cholerae* FIGR recipients. One-way ANOVA with Dunnett’s multiple comparison test comparing each strain to the ‘Cre only’ strain. *, p<0.05; **, p<0.01; ***, p<0.001; ns, not significant

**Supplemental Figure S3:**
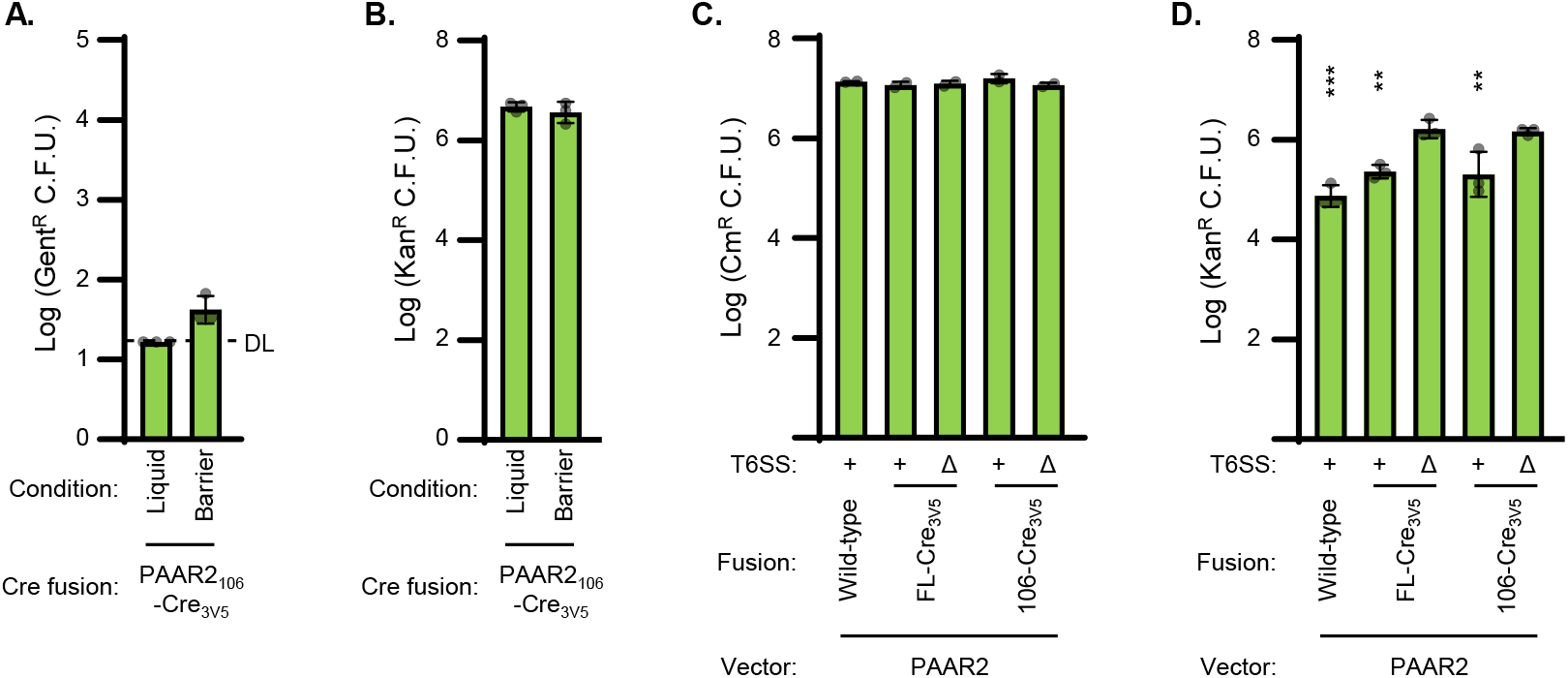
PAAR2 fusions total donor and recipient survival, liquid or barrier controls, related to Figure 2B. A, B. Cre delivery in liquid suspension (liquid) or when donor and recipient were separated by a nitrocellulose membrane (barrier). Recovery of recipient colony forming units after incubating with wild-type *V. cholerae* with PAAR2_106_-Cre_3V5_. **A.** Gent^R^ C.F.U. indicative of Cre-mediated FIGR cassette recombination. DL, detection limit. **B.** Total recipient C.F.U. (Kan^R^). **C.** Recovery of *V. cholerae* Cm^R^ colony forming units (total donor cell recovery) with indicated Cre fusions to PAAR2 after incubation with *V. cholerae* FIGR recipients. One-way ANOVA analysis indicates no significant differences. **D.** Recovery of *V. cholerae* Kan^R^ (total recipients) colony forming units after delivery from active T6SS (+) or Δ*tssM* (Δ) *V. cholerae* with indicated Cre fusions to PAAR2 vectors. One-way ANOVA with Sidak’s multiple comparison test comparing active T6SS and Δ*tssM* donors with comparable delivery fusions. **, p<0.01; ***, p<0.001.

**Supplemental Figure S4:**
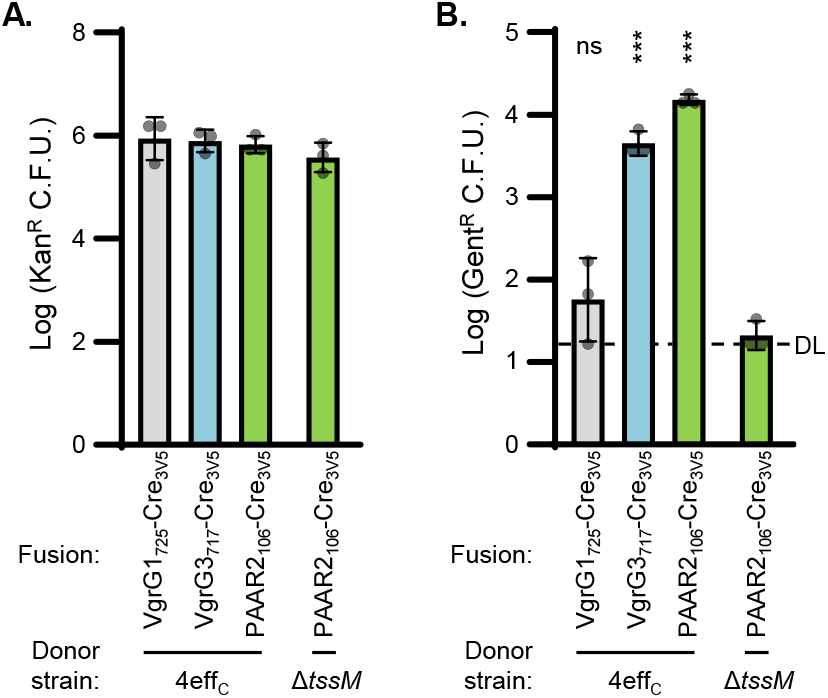
Cre fusion delivery from effectorless donor strain, related to Figure 2C. **A.** Total recipients recovered (Kan^R^ C.F.U.) after delivery from *V. cholerae* with catalytically inactivated antibacterial effectors (4eff_C_) or Δ*tssM* (Δ). One-way ANOVA with Dunnett’s multiple comparison test comparing each sample to the Δ*tssM* donor strain. ***, p<0.001; ns, not significant. **B.** As for **A.** but showing Cre-recombined recipient bacteria (Gent^R^ C.F.U). DL, detection limit. One-way ANOVA analysis indicates no significant differences.

**Supplemental Figure S5:**
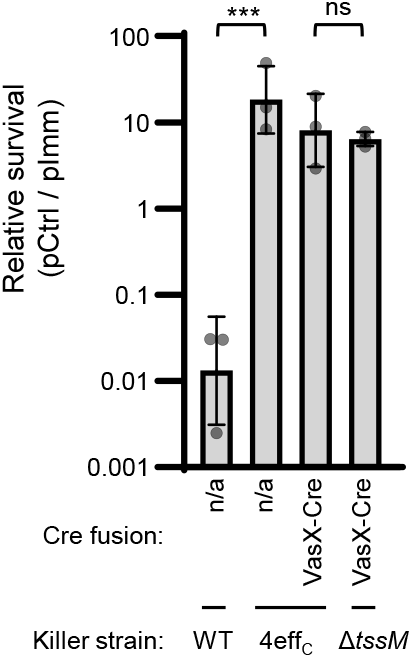
The VasX-Cre fusion does not kill sensitive prey. T6SS competition assay showing killing by V52 wild-type (WT), catalytically inactivated antibacterial effectors (4eff_C_), or inactivated T6SS (Δ*tssM*). Strains were complemented with no plasmid (n/a), or VasX-Cre. Prey are a V52 strain lacking the VasX immunity gene, *tsiV2*, and complemented with either *tsiV2* (pImm) or a non-protective chaperone gene as a control (VCA0019; pCtrl). Survival of the prey is shown as a ratio of recovered C.F.U. (pCtrl / pImm). One-way ANOVA with Sidak’s multiple comparison test comparing no plasmid WT or VasX-Cre delivery to controls. ***, p<0.001; ns, not significant.

**Supplemental Figure S6:**
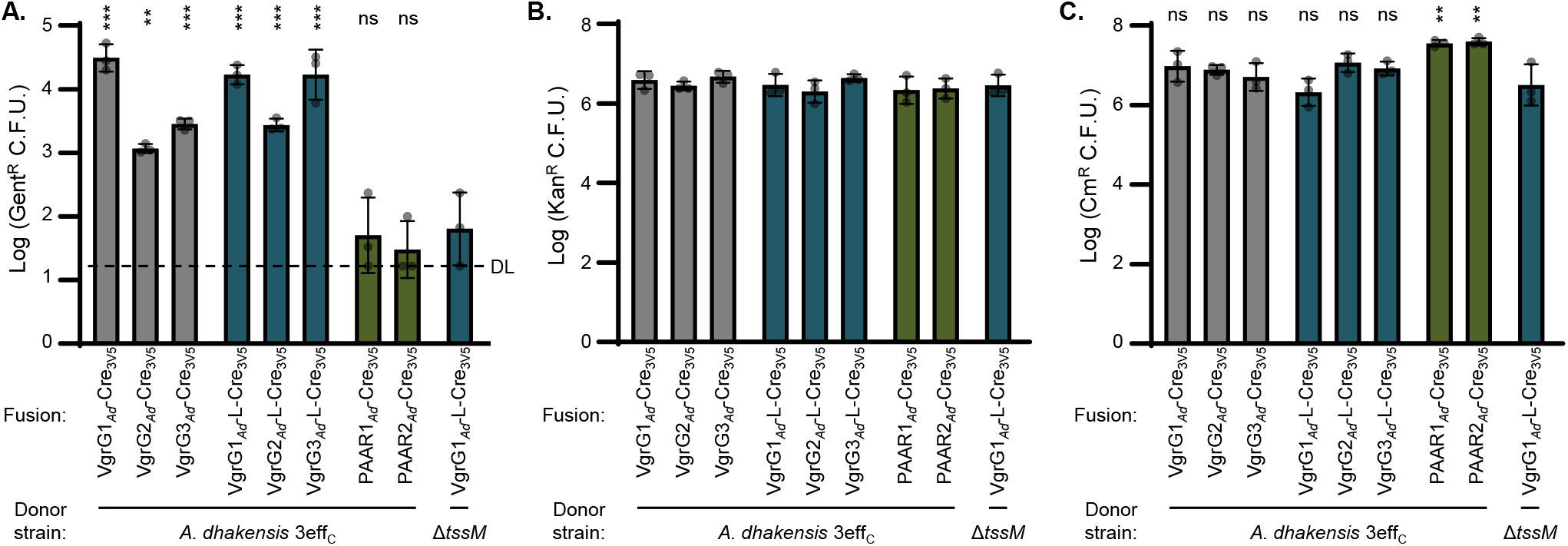
*A. dhakensis* donor total recipient survival and recombinant recovery, related to Figure 3C. A. Recovery of Cre-recombined recipient bacteria (Gent^R^ C.F.U.) after delivery of Cre fusions from *A. dhakensis* with catalytically inactivated antibacterial effectors (3eff_C_) or Δ*tssM* (Δ). DL, detection limit. One-way ANOVA with Dunnett’s multiple comparison test comparing each sample to the Δ*tssM* donor. **, p<0.01; ***, p<0.001; ns, not significant. **B.** As for **A.** but showing total recipients recovered (Kan^R^ C.F.U.). One-way ANOVA analysis indicates no significant differences. **C.** Recovery of *A. dhakensis* CmR colony forming units (total donor cell recovery) with indicated Cre fusions after incubation with *V. cholerae* FIGR recipients. One-way ANOVA with Dunnett’s multiple comparison test comparing each sample to the Δ*tssM* donor. **, p<0.01; ns, not significant.

**Supplemental Figure S7:**
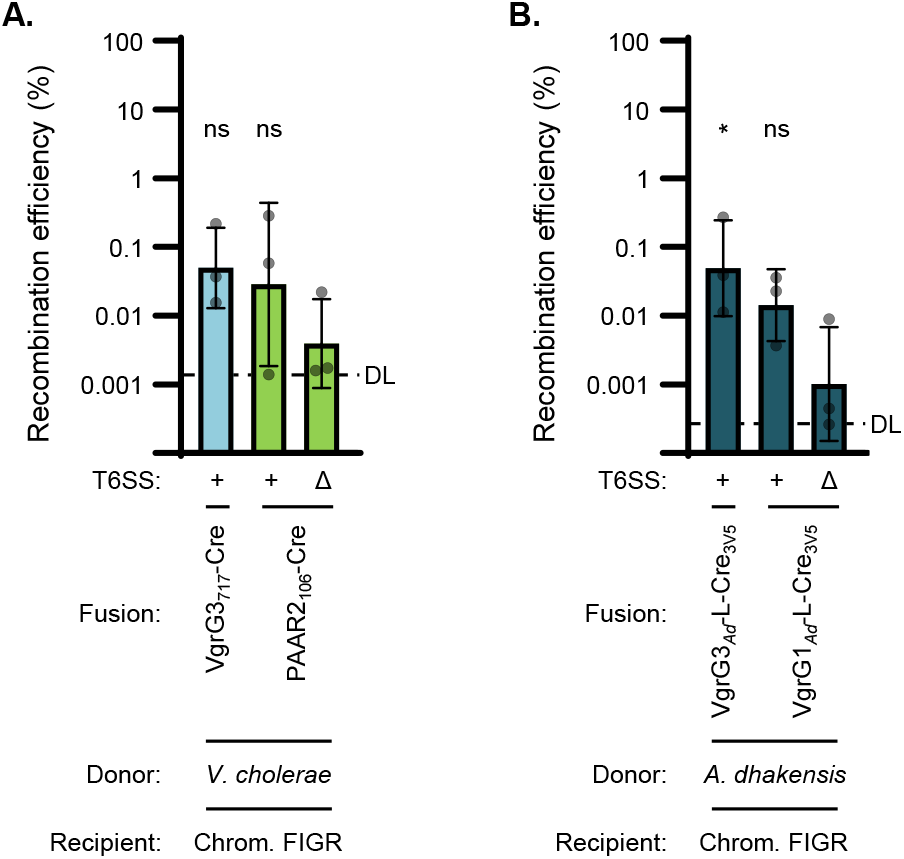
Cre delivery to *V. cholerae* recipients with chromosomal FIGR. **A.** Recombination efficiency of *V.cholerae* recipients with chromosomal FIGR cassette after delivery from *V. cholerae* 4eff_C_ (+) or Δ*tssM* (Δ) strains. **B.** Recombination efficiency of *V.cholerae* recipients with chromosomal FIGR cassette after delivery from *A. dhakensis 3*eff_C_ (+) or Δ*tssM* (Δ) strains. DL, approximate detection limit. For each experiment, one-way ANOVAs with Dunnett’s multiple comparison tests comparing samples to Δ*tssM* donors. *, p<0.05; ns, not significant.

**Supplemental Table S1:Strains and plasmids used in this study**

